# A 6-minute Limb Function Assessment for Therapeutic Testing in Experimental Peripheral Artery Disease Models

**DOI:** 10.1101/2024.03.21.586197

**Authors:** Victoria R. Palzkill, Jianna Tan, Abigail L. Tice, Leonardo F. Ferriera, Terence E. Ryan

## Abstract

**Background:** The translation of promising therapies from pre-clinical models of hindlimb ischemia (HLI) to patients with peripheral artery disease (PAD) has been inadequate. While this failure is multifactorial, primary outcome measures in preclinical HLI models and clinical trials involving patients with PAD are not aligned well. For example, laser Doppler perfusion recovery measured under resting conditions is the most used outcome in HLI studies, whereas clinical trials involving patients with PAD primarily assess walking performance. Here, we sought to develop a 6-min limb function test for preclinical HLI models that assess muscular performance and hemodynamics congruently.

**Methods:** We developed an *in situ* 6-min limb function test that involves repeated isotonic (shortening) contractions performed against a submaximal load. Continuous measurement of muscle blood flow was performed using laser Doppler flowmetry. Quantification of muscle power, work, and perfusion are obtained across the test. To assess the efficacy of this test, we performed HLI via femoral artery ligation on several mouse strains: C57BL6J, BALBc/J, and MCK-PGC1α (muscle-specific overexpression of PGC1α). Additional experiments were performed using an exercise intervention (voluntary wheel running) following HLI.

**Results:** The 6-min limb function test was successful at detecting differences in limb function of C57BL6/J and BALBc/J mice subjected to HLI with effect sizes superior to laser Doppler perfusion recovery. C57BL6/J mice randomized to exercise therapy following HLI had smaller decline in muscle power, greater hyperemia, and performed more work across the 6-min limb function test compared to non-exercise controls with HLI. Mice with muscle-specific overexpression of PGC1α had no differences in perfusion recovery in resting conditions, but exhibited greater capillary density, increased muscle mass and absolute force levels, and performed more work across the 6-min limb function test compared to their wildtype littermates without the transgene.

**Conclusion:** These results demonstrate the efficacy of the 6-min limb function test to detect differences in the response to HLI across several interventions including where traditional perfusion recovery, capillary density, and muscle strength measures were unable to detect therapeutic differences.

## INTRODUCTION

Lower extremity peripheral artery disease (PAD) is caused by decreased blood flow to the leg resulting from atherosclerosis. PAD is estimated to affect more than 230 million people worldwide and increases the risk for other cardiovascular events (coronary heart disease and stroke) and limb amputation (1). Current medical management for PAD revolves heavily around reducing the risk of cardiovascular events and slowing PAD progression, or for surgical revascularization for patients with severe PAD. Unfortunately for patients with PAD, therapeutic progress for improving lower limb function has been stagnant for decades. Promising results on cell- and gene-based angiogenesis therapies in pre-clinical (animal) models of PAD led to several clinical trials in patients with PAD (2–13). Sadly, these interventions have failed to reach the clinic to date. As a result, cilostazol (a PDE3 inhibitor) remains the only pharmacological therapy to improve walking performance in patients with PAD (pentoxifylline is approved by the Food and Drug Administration in the USA but its performance is inferior to cilostazol, and thus clinical usage is limited).

While the failure of therapeutic angiogenesis trials is multifactorial, there is a major gap in the methodology used to assess therapeutic efficacy with pre-clinical models of PAD and those used to assess clinical efficacy in randomized trials involving patients with PAD. The most commonly used pre-clinical model of PAD is hindlimb ischemia (HLI), induced by ligation and/or excision of the femoral artery of the hindlimb of animals including rodents (14) and rabbits (15, 16). Variations of this procedure have been developed to modulate the severity and longevity of the ischemic condition within the surgical limb (17–21). The overwhelming majority of pre-clinical HLI studies employ laser Doppler to quantify limb perfusion recovery in anesthetized animals (22) as a primary outcome measure, which is often supported with histochemical and/or imaging-based analyses of vascularity. In contrast, assessments of walking performance have emerged as the gold standard primary outcome measure in clinical trials involving patients with PAD (13, 23–26). Increasingly, the 6-minute walk test has been preferred over treadmill walking tests (24, 27). Walking performance analysis can encompass multiple domains of PAD pathophysiology including decrease oxygen delivery, impaired oxygen utilization (mitochondrial dysfunction), and poor muscle strength/power or motor coordination. However, assessing walking performance in pre-clinical HLI models is challenging as quadrupedal animals can have normal walking speeds despite the surgical induction of HLI in a single hindlimb. To address this limitation, we developed a novel 6-minute limb function test that utilizes electrical stimulation to quantify muscle perfusion, power, and total work simultaneously. We assessed the ability of this test to distinguish between mouse strains with known differences in HLI responses, HLI mice enrolled in an exercise training intervention, and transgenic mice that overexpress peroxisome proliferator-activated receptor gamma coactivator 1-alpha, a known regulator of muscle mitochondrial biogenesis and angiogenesis, specifically in striated muscle (MCK-PGC1α).

## MATERIALS AND METHODS

### Animals

Experiments were conducted on male 12-week-old BALB/cJ (Stock No. 000651) and C57BL6J (Stock No. 000664) mice purchased from Jackson Laboratories. Mice with transgenic overexpression of peroxisome proliferator-activated receptor gamma coactivator 1-alpha (*Ppargc1a*, also known as PGC1α) under the direction of the muscle creatine kinase promoter (C57BL/6-Tg(Ckm-Ppargc1a)31Brsp/J; Stock No. 00823) (28) were obtain from Jackson Laboratories. Hemizygous transgenic mice (termed MCK-PGC1α herein) were bred with non-carrier C57BL6J mice (Stock No. 000664) and wildtype littermates without the transgene were used as controls. Both the researchers and surgeon were blinded to the genotype and/or group of the animals. All mice were housed in temperature (22°C) and light-controlled (12:12-h light-dark) rooms and maintained on standard chow (Envigo Teklad Global 18% Protein Rodent Diet 2918 irradiated pellet) with free access to food and water prior to enrollment. All animal experiments adhered to the *Guide for the Care and Use of Laboratory Animals* from the Institute for Laboratory Animal Research, National Research Council, Washington, D.C., National Academy Press. All procedures were approved by the Institutional Animal Care and Use Committee of the University of Florida (Protocols 202110484 and 202010121).

### Animal model of peripheral artery disease

Femoral artery ligation (29, 30) was performed by anesthetizing mice with intraperitoneal injection of ketamine (100 mg/kg) and xylazine (10 mg/kg) and surgically inducing unilateral hindlimb ischemia (HLI) by placing silk ligatures on the femoral artery just distal the inguinal ligament and immediately proximal to the saphenous and popliteal branches. Extended-release buprenorphine (3.25 mg/kg) was given post-operatively for analgesia.

### Voluntary wheel running exercise

Mice randomized to the exercise training group were placed into cages contain a running wheel (Actimetrics) immediately following surgical induction of HLI. Mice were housed individually in the wheel cages for 21-days post-HLI to allow for quantification of running distance for each mouse. Daily voluntary running distance was measured using ClockLab analysis software (Actimetrics).

### Limb perfusion assessment

Limb perfusion was assessed by laser Doppler flowmetry (moorVMS-LDF, Moor Instruments) prior to surgery, immediately post-surgery, and every seven days post-surgery until euthanasia as described previously (29, 31, 32). Both hindlimbs were shaved and the laser Doppler probe was placed ∼1-2mm away from the middle of the posterior side of the paw, the posterior side of the lateral head of the gastrocnemius muscle, and the midbelly of the tibialis anterior. Perfusion recovery was reported as a percentage of the non-ischemic limb.

### Nerve mediated isometric muscle contraction

Functional tests of the plantar flexor muscles, specifically the gastrocnemius/soleus/plantaris (GPS) complex was measured *in-situ* using a whole animal system (model 1300A, Aurora Scientific Inc. Aurora, ON, Canada). Mice were anaesthetized with an intraperitoneal injection of xylazine (10 mg/kg) and ketamine (100 mg/kg) and subsequent doses were given as needed for maintenance. The GPS complex from the ischemic limb was isolated from its distal insertion leaving all vasculature intact. The distal portion of calcaneal tendon was tied using a 4-0 silk suture attached to the lever arm of the force transducer (Cambridge Technology; Model No. 2250). Muscle contractions were elicited by stimulating the sciatic nerve via bipolar electrodes using square wave pulses (Aurora Scientific, Model 701A stimulator). Lab-View-based DMC program (version 615A.v6.0, Aurora Scientific Inc.) was used for data collection and servomotor control. After obtaining optimal length via twitch contractions, an abbreviated force frequency curve was performed. Isometric contractions were elicited using 500ms train (current 2mA, pulse width 0.2ms) at stimulation frequencies of 1Hz, 40Hz, 80Hz, and 150Hz with one minute of rest between contractions. The peak tetanic force generated from the 80Hz contraction was used as the reference force for subsequent isotonic contractions. Tetanic force levels were reported as absolute force and specific force (absolute force normalized to muscle mass).

### 6-min Limb Function Test

To evaluate muscular performance across a series of repetitive contractions, we chose to utilize afterloaded isotonic contractions which involve electrically stimulating the nerve with supramaximal voltage (2mA, 0.2ms pulse width, 100ms train duration) at 80Hz to activate all motor units and allowing the muscle to shorten once it reaches the prescribed isometric tension (30% or 45% of the 80Hz peak isometric force). This contraction pattern is more similar to voluntary muscle activation and allows for the quantification several muscle performance characteristics including force (newtons), displacement (mm), shortening velocity (m/s), mechanical power (watts), and mechanical work (joules). We calculated muscle shortening velocity as the change in distance (mm) from a 10ms period which began 20ms after the initial length change. Mechanical peak power was calculated as the product of the shortening velocity (m/s) and corresponding force (N/kg). We also calculated instantaneous velocity as the first derivative of the position-time tracing (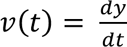; where *y* is position and *t* is time). From this, we calculated instantaneous power as the product of the instantaneous velocity and the corresponding force (N/kg). Within each contraction, we then quantified mechanical work as the integral of the instantaneous power-time tracing (Work = ∫ *Instantaneous Power* · *dt*; where *t* is time).

To assess limb function in mice with HLI, we designed a sequence in which the afterloaded isotonic contraction described above was followed by a 2.5 second static rest period to allow Laser Doppler perfusion measurement, and then three sinusoidal 0.1mm passive stretches from optimal length to stabilize the sarcomere (33–36). This contraction, rest (perfusion assessment), passive stretch sequence was repeated 86 times for a total test duration of 6 minutes. Laser Doppler flowmetry was employed by placing the probe on the medial gastrocnemius throughout the 6-min test to assess functional hemodynamics. Average perfusion flux values during the 2.5ms static rest period immediately following each contraction. The degree of muscular fatigue was presented as the change in power (watts/Kg) and as a percentage of the initial power across the 6-min test. Mechanical isotonic work was also quantified for each contraction and changes in work were plotted across the 6-min test. The sum of work performed across the 6-min test was also calculated. All muscle testing was performed on a temperature-controlled platform to maintain a body temperature of 37°C.

### Labeling of Perfused Capillaries

Mice received a retro-orbital injection of 50 µL of 1 mg/mL Griffonia simplicifolia lectin (GSL) isolectin B4, Dylight 649 (Vector Laboratories; Cat. No. DL-1208) to fluorescently label α-galactose residues on the surface of endothelial cells of perfused capillaries. Following the injection, animals were returned to their cage and allowed 1-2 hours of free movement prior to the 6-min limb function test and subsequent euthanasia and tissue harvesting.

### Immunofluorescence microscopy

Following completion of the 6-min limb function test, the plantarflexor complex (gastrocnemius, soleus, plantaris muscles) were carefully dissected, weighed, embedded in optimal cutting temperature (OCT) compound, and rapidly frozen in liquid nitrogen-cooled isopentane for cryosectioning. Using a Leica 3050S cryotome, 10µm-thick transverse sections of the plantarflexors were cut and mounted on microscope slides. All muscle sections were fixed with 4% paraformaldehyde (ThermoFisher Scientific, Cat. No. J19943-K2) for ten minutes, and permeabilized with 0.25% triton X-100 (Millipore-Sigma, Cat. No. 93443). Following three washes with 1X phosphate buffered saline (PBS), slides were incubated in blocking solution (5% goat serum + 1% BSA in 1XPBS) for 4-6 hours. To label total capillaries, muscles were incubated with a primary antibody against PECAM1 (anti-CD31, Abcam, Cat. No. ab28364, 1:100 dilution) overnight at 4°C. The following day slides were washed with 1XPBS and incubated for one hour at room temperature with Alexa-Fluor555 anti-rabbit secondary antibody (ThermoFisher Scientific, Cat. No. A11034, 1:250 dilution). On a separate slide, muscle sections were incubated overnight at 4°C with a primary antibody against laminin (Millipore-Sigma, Cat. No. L9393, 1:100 dilution) to label the basal lamina surrounding myofibers, and alpha-smooth muscle actin (ThermoFisher Scientific, Cat. No. 14976082; 1:400 dilution) to label arterioles. The sections were then washed with 1xPBS and stained with Alexa Fluor 555 goat anti-mouse IgM (ThermoFisher Scientific, Cat. No. A-21426, 1:250 dilution) and Alexa Fluor 488 goat anti-rabbit IgG (ThermoFisher Scientific, Cat. No. A-21121, 1:250 dilution) secondary antibodies. Coverslips were mounted onto all slides using Vectashield hardmount (Vector Laboratories, Cat. No. H-1500). Slides were imaged at 20x magnification with an Evos FL2 Auto microscope (ThermoFisher Scientific), and tiled images of the entire section were obtained for analysis. Images were thresholded and the amount of perfused capillaries, total capillaries, and arterioles were quantified by a blinded investigator using a particle counter in Fiji/ImageJ. Skeletal myofiber cross-sectional area (CSA) was quantified using MuscleJ2 (37), an automated analysis software developed in ImageJ/Fiji.

### Study Approval

All procedures were approved by the Institutional Animal Care and Use Committee of the University of Florida (Protocols 202110484 and 202010121).

### Data Availability

All source data are available from

### Statistical Analysis

All data are presented as the mean ± standard error (SEM). Normality of data was tested with the Shapiro-Wilk test and/or inspection of QQ plots. Data involving comparisons of two groups were analyzed using a student’s t-test when normally distributed and Mann-Whitney test when normality could not be assessed. When making comparisons with repeated measures, data were analyzed using repeated measures ANOVA with Tukey’s post hoc testing for multiple comparisons when significant interactions were detected. In all cases, *P* < 0.05 was considered statistically significant. All statistical testing was conducted using GraphPad Prism software (version 9.0).

## RESULTS

### Developing a 6-minute limb function test to evaluate isotonic muscle function and perfusion

Our first objective was to design a hindlimb function testing protocol that facilitates analysis of both muscular performance (e.g., power or work) and vascular hemodynamics (e.g., blood flow), two factors that are important for walking performance in patients with PAD. To accomplish this, we adapted methods used in our previous studies to evaluate muscle contraction and limb perfusion (31, 38–40). In this regard, we chose to develop the test with a focus on the plantarflexor muscles, which are important for both mouse and human locomotion, affected by surgical HLI in mice, and can be stimulated via the sciatic nerve in situ where the muscles are perfused by the native vasculature. In contrast to our previous work using isometric contractions where the tendon-tendon length remains constant, we employed an isotonic contraction protocol where the muscle is allowed to shorten against a submaximal load. We chose a submaximal stimulation frequency of 80Hz, which produced a fused tetanic contraction, as our reference force (**Figure 1A**). With the isotonic contraction, the lever arm is allowed to shorten once the muscle exceeds the desired reference force is reach during a contraction. We tested load clamps at both 30% and 45% of the reference force (peak force of the 80Hz isometric contraction) as shown in **Figure 1B**. As expected, the higher percentage of the reference force (45%) resulted in slower and less movement of the level arm in the position-time tracing (**Figure 1B**) – consistent with the strain dependence of the acto-myosin crossbridge (41). Because the load/force is constant and both distance and time can be measured, the muscle shortening velocity (distance/time), power (force x velocity), and work (integral of power-time) can be quantified for each contraction (**Figure 1C**). This contraction protocol was chosen for several reasons: 1) the isotonic contraction is more closely relevant to the contractions the plantarflexor muscles experience during locomotion; 2) the movement during the isotonic contraction allows quantification of total work (integral of power x time) and outcome akin to walking distance in patients; and 3) muscle perfusion can be assessed between contractions to assess the hyperemic response (**Figure 1D**). Following the completion of a contraction, a ∼2.5 second period of rest was provided to allow analysis of limb perfusion (without motion artefact) via laser Doppler flowmetry (**Figure 1E**). Finally, three brief passive stretches (0.1mm) were used to realign sarcomere lengths (**Figure 1E**). This sequence was repeated 86 times (once every ∼4.2s) to produce a 6-minute limb function test.

**Figure 1.**
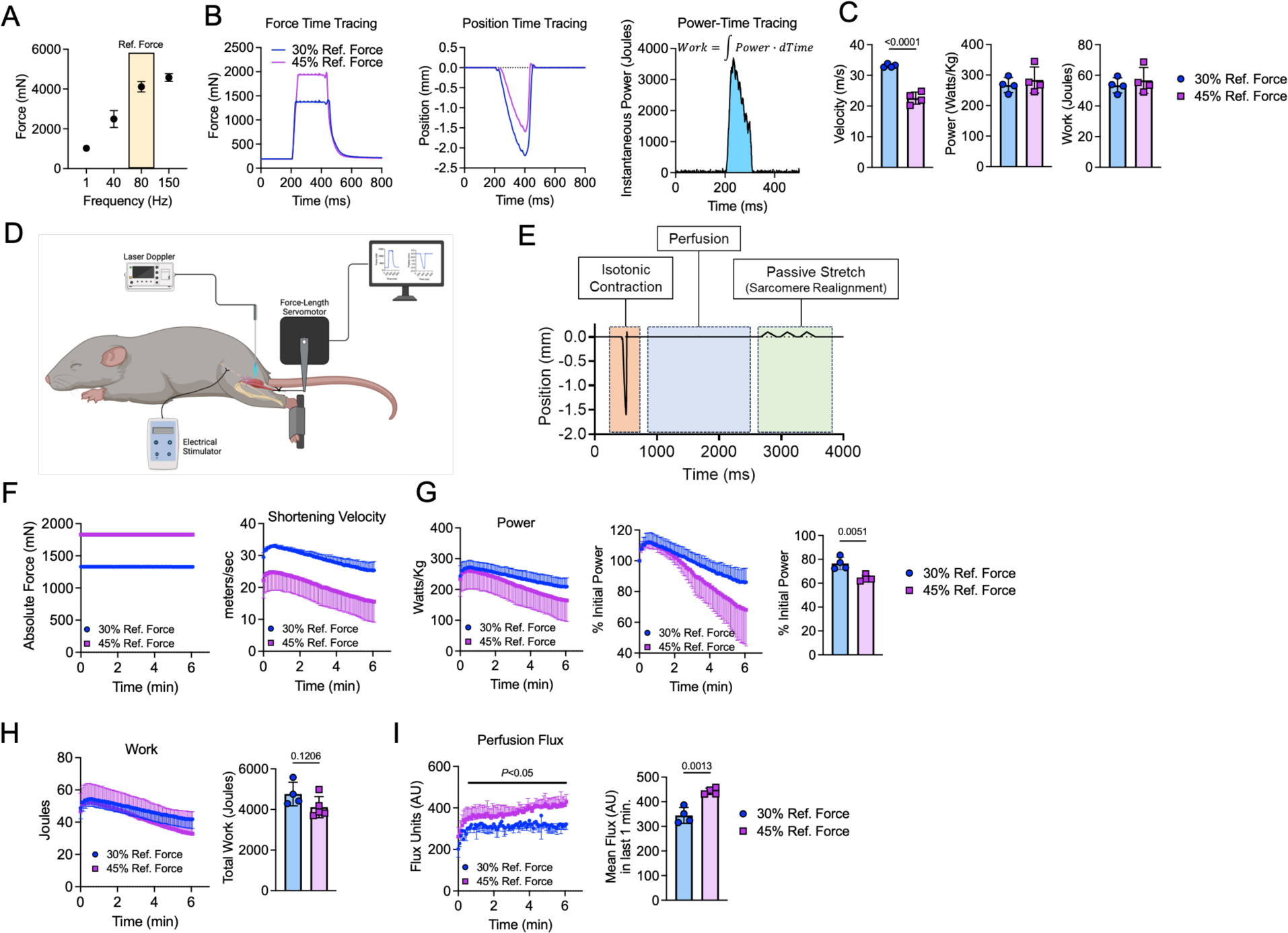
Developing a 6-minute limb function test to evaluate isotonic muscle function and perfusion. (A) Force-frequency curve of the gastrocnemius muscle in healthy mice (n=8) showing selection of the 80Hz contraction as reference force. (B) Force-time and position-time tracings showing levels of force production and length change (shortening) at 30% and 45% of the reference force. Calculation of the Power-Time tracing allows for quantification of work. (C) Quantification of shortening velocity, power, and work (n=4/group). (D) Graphic of the experiment setup. (E) Graphical representation of the isotonic contraction protocol used in the 6-min test. (F) Force and shortening velocity across the 6-min test (n=4/group). (G) Power expressed both in absolute units (watts) and a percentage of the initial power across the 6-min test and quantification of the percent power loss at the end of the test (n=4/group). (H) Work performed across the 6-min test and the total work summed (n=4/group). (I) Laser Dopper perfusion flux across the 6-min test and the mean flux quantified during the last one-minute (n=4/group) Panel C was analyzed using unpaired, two-tailed Student’s *t*-test. Data were analyzed using an unpaired *t*-test (two-tailed). Error bars represent the standard error. The graphic and panel D was generated using BioRender.

Using healthy mice without HLI, we first aimed to validate that the testing protocol could detect differences in muscle performance and perfusion when performed at different loads (30% vs. 45% of the reference force). As expected, the force of the 45% group was higher than the 30% group, but forces in both groups was unchanged across the 6-min test (**Figure 1F**). As expected, shortening velocity was higher in the 30% group because the force is lower when compared to the 45% group (**Figure 1F**). Regardless of group, shortening velocity decreased across the 6-min test which is indicative of muscle fatigue. Muscle peak power, expressed as either watts/kg or a percentage of the initial power, declined across the 6-min test (**Figure 1G**) and the percent power loss at the end of the test was greater in the 45% group (*P* = 0.0051). Total work performed was similar between groups (**Figure 1H**), however the 45% group had significantly higher gastrocnemius perfusion rates (**Figure 1I**). These data demonstrate the ability to distinguish dose-dependent differences in muscle performance and hemodynamics using this 6-min test.

### 6-min Limb Function Testing Distinguishes Strains with Differing Sensitivity to HLI

As the next step in evaluating the efficacy of the 6-min limb function test in HLI, we leveraged known genetic differences in the ischemia-resistant C57BL6/J and ischemia-sensitive BALBc/J mouse strains (18, 19, 42–49). Consistent with the large body of evidence, BALB/cJ mice displayed lower paw and gastrocnemius perfusion recovery at rest (**Figure 2A**) as well as fewer total and perfused capillaries (**Figure 2B**) and arterioles (**Figure 2C**) when compared to C57BL6/J mice subjected to HLI. Similarly, the gastrocnemius mass (**Figure 2D**) and mean myofiber cross-sectional area (CSA) (**Figure 2E**) were also significantly lower in BALBc/J mice. Isometric forces, presented as both absolute and specific (normalized to muscle mass), were significantly lower in BALBc/J mice with HLI compared to C57BL6/J mice with HLI (**Figure 2F**). A representative position-time tracing demonstrates the marked decrease in the ability of the BALBc/J mice to move the lever arm (**Figure 2G**), and consequently peak power was reduced by ∼80% in the BALBc/J mice (**Figure 2G**). Muscle power changes across the 6-min test are shown in **Figure 2H** as both Watts/Kg and the percentage of the initial power. Muscle power was substantially higher in C57BL6/J mice compared to BALBc/J mice across the entire testing period, however, the percent decline in muscle power was greater in C57BL6/J mice (**Figure 2H**). C57BL6/J mice performed substantially more work across the test when compared to BALBc/J mice with HLI (**Figure 2I**), demonstrating that the capacity for limb work is significantly better in C57BL6/J mice. Consistent with the C57BL6/J genetic advantage of having higher collateral vessel density (49–51), the hyperemic response in gastrocnemius perfusion flux was significantly greater in C57BL6/J (**Figure 2J**).

**Figure 2.**
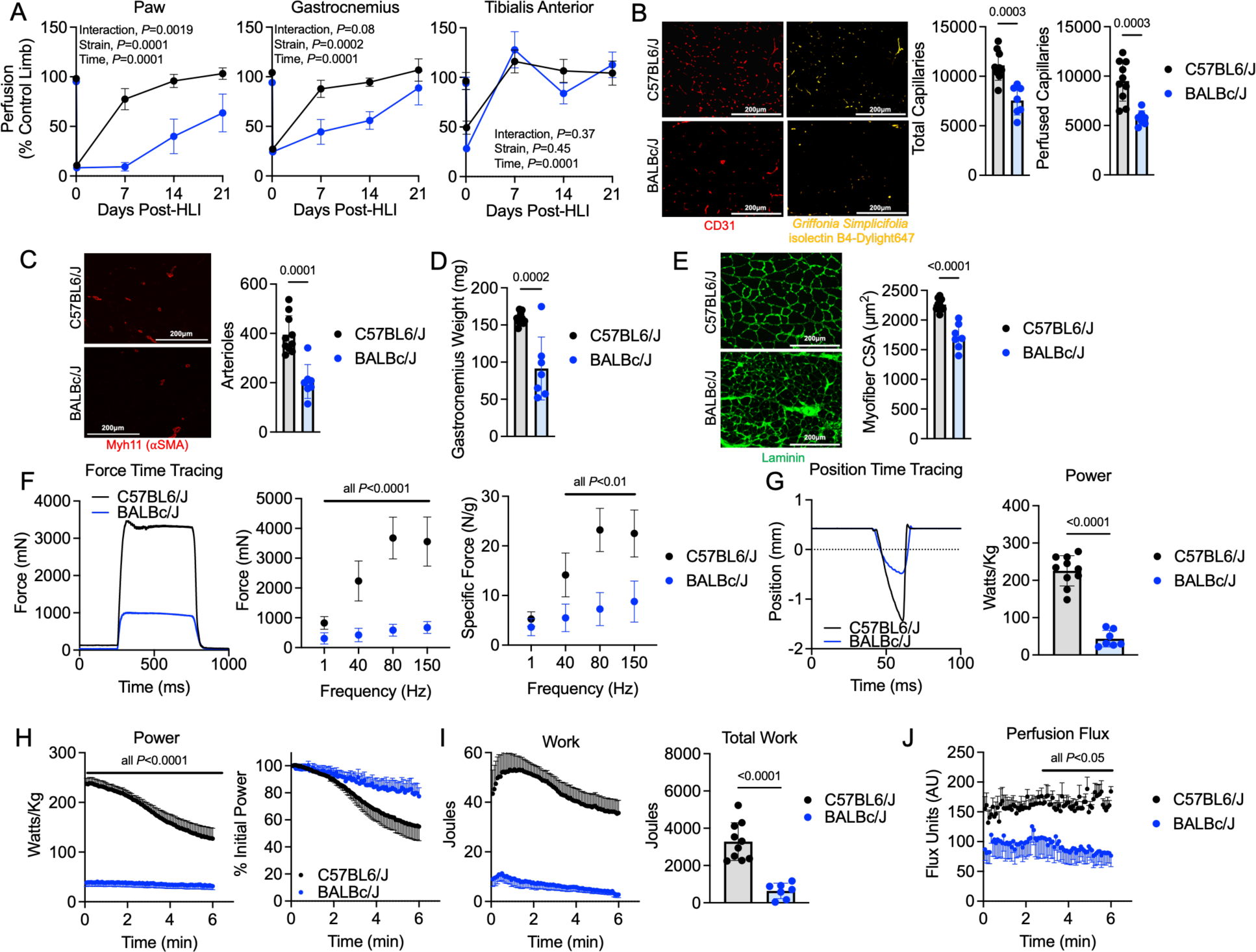
6-min Limb Function Testing Distinguishes Strains with Differing Sensitivity to HLI. (A) Laser Doppler flowmetry quantification of perfusion recovery in the paw, gastrocnemius, and tibialis anterior muscles expressed as percentage of control limb (n=7-10/group). Perfusion recovery was analyzed using two-way ANOVA. (B) Representative immunofluorescence images and quantification of total and perfused capillaries. (C) Representative immunofluorescence images and quantification of arterioles. (D) Gastrocnemius muscle mass. (E) Representative images and quantification of the myofiber cross-sectional area (CSA). (F) Representative force traces of 80Hz contractions between strains as well as quantification of absolute and specific forces at different stimulation frequencies. (G) A representative position-time tracing and quantification of muscle power. (H) Muscle power across the 6-min test. (I) Work performed across the 6-min test and quantification of the total work completed in the test. (J) Perfusion flux during the 6-min test. Statistical analyses in panels B-J were performed using an unpaired *t*-test (two-tailed). Error bars represent the standard error.

### Exercise Training Improves Limb Performance in Mice with Experimental Peripheral Artery Disease

Supervised exercise training has emerged as one of the most effective treatments to improve or prevent declines in walking performance in patients with PAD (1, 13, 23, 52–54). Additional evidence has begun to show that home-based exercise interventions can also improve walking performance in patients with PAD (26, 55), Based on these clinical observations, we next sought to determine if this 6-min limb function test could distinguish differences in mice subjected to HLI and randomized to either control or exercise training for three weeks (**Figure 3A**). Exercise training consisted of voluntary wheel running in which C57BL6/J mice were placed in wheel cages immediately following HLI surgery, whereas the control group was housed in cages without running wheels. The average running distance increased across the post-HLI period with mice reaching ∼8Km/d in the days immediately before euthanasia (**Figure 3B**). Laser Doppler perfusion recovery, measured in the resting condition of anesthetized mice, was not different between control and exercise groups for either the paw, gastrocnemius, or tibialis anterior muscles (**Figure 3C**). Consistent with the wealth of literature, exercise trained mice had higher total capillary densities compared to control mice with HLI (*P* =0.018), however the perfused capillary density was not different (**Figure 3D**). There was a non-significant trend (*P* = 0.0689) for increased arteriole density in the exercise group (**Figure 3E**). There were no differences in the mean myofiber CSA (**Figure 3F**) or absolute isometric force production (**Figure 3G**). However, there was a modest, but significant increase in gastrocnemius mass in the exercise group (**Figure 3H**), but this did not result in significant changes in isometric specific force levels (**Figure 3I**). Peak power, measured using the initial velocity in the first 15ms of the shortening contraction, were not different between groups (**Figure 3J**). However, the total distance change, as seen in the position-time tracing, was greater in mice randomized to the exercise group post-HLI (**Figure 3J**). While both control and exercise mice had similar muscle power at the start of the 6-min test, exercise trained mice with HLI experienced less decline in muscle power than control mice with HLI (**Figure 3K**). Quantification of work using the integral of the instantaneous power and time, fully demonstrated the superior limb function in the exercise mice (**Figure 3L**), a result akin to the increase in walking performance observed in patients with PAD following supervised or home-based exercise training. Another important observation from the 6-min test was that exercise mice had significantly higher gastrocnemius perfusion flux during the test when compared to control mice (**Figure 3M**). This is in contrast to the perfusion recovery assessments performed in resting conditions where control and exercise mice were not different (**Figure 3C**). This discrepancy highlights the value of assessing limb hemodynamics under conditions where muscle contraction facilitates the ability to assess the increase limb/muscle perfusion with muscular work.

**Figure 3.**
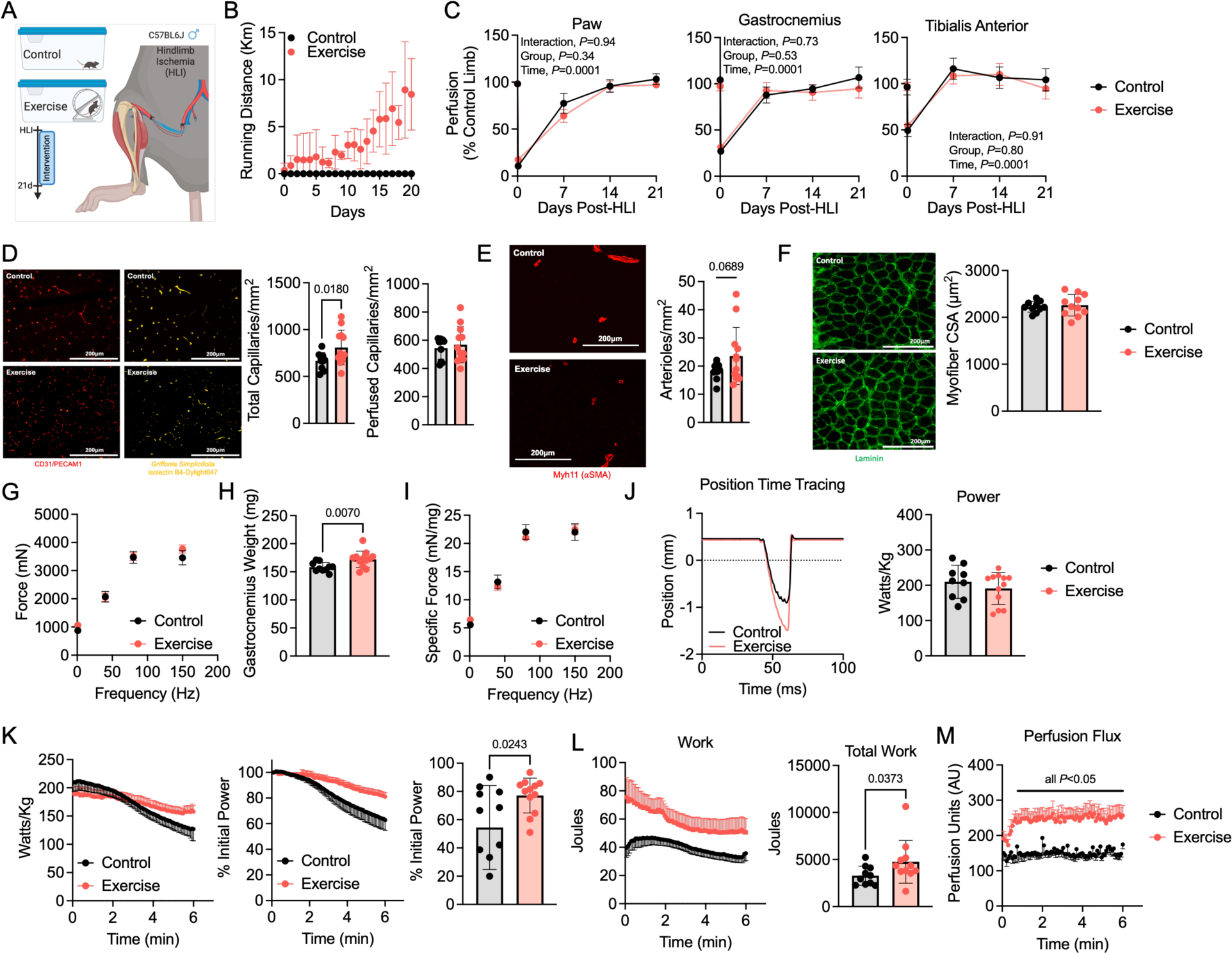
Exercise Training Improves Limb Performance in Mice with Experimental Peripheral Artery Disease. (A) Graphic of the experimental design and timeline (generated using BioRender). (B) Daily running distance across the intervention. (C) Laser Doppler flowmetry quantification of perfusion recovery in the paw, gastrocnemius, and tibialis anterior muscles expressed as percentage of control limb (n=10-12/group). Perfusion recovery was analyzed using two-way ANOVA. (D) Representative immunofluorescence images and quantification of total and perfused capillaries. (E) Representative immunofluorescence images and quantification of arterioles. (F) Representative images and quantification of the myofiber cross-sectional area (CSA). (G) Quantification of absolute forces at different stimulation frequencies. (H) Gastrocnemius muscle mass. (I) Quantification of specific forces at different stimulation frequencies. (J) A representative position-time tracing and quantification of muscle power. (K) Muscle power across the 6-min test and quantification of power loss at the end of the test. (L) Work performed across the 6-min test and quantification of the total work completed in the test. (M) Perfusion flux during the 6-min test. Panels D-M were analyzed using an unpaired *t*-test (two-tailed). Error bars represent the standard error. All panels contain n=10-12/group.

### Muscle-specific Overexpression of PGC1α Improves Limb Function Following HLI

Mitochondrial impairments in skeletal muscle of patients with PAD have been well-documented in the literature (56–72). It is believed that compromised oxygen utilization secondary to pathological changes in mitochondrial function may contribute to impaired walking performance and limb function. To explore this possibility using the 6-min limb function test, we performed HLI on transgenic mice that overexpress PGC1α, a known regulator of mitochondrial biogenesis muscle adaptation (28, 73–79), under the control of the muscle creatine kinase promoter (termed MCK-PGC1α) and their wildtype (WT) littermates (**Figure 4A**). Laser Doppler perfusion recovery, measured under the resting condition, was not different between MCK-PGC1α mice and their wildtype (WT) littermates following HLI (**Figure 4B**). Consistent with previous reports (79, 80), MCK-PGC1α mice had higher total and perfused capillary densities compared to WT mice (**Figure 4C**). However, arteriole density was not different between MCK-PGC1α and WT mice (**Figure 4D**). Absolute isometric forces were significantly higher in MCK-PGC1α compared to WT mice at higher stimulation frequencies (**Figure 4E**). The increase muscle strength was primarily driven by increase muscle mass in MCK-PGC1α mice (**Figure 4F**) as specific forces were not different between groups (**Figure 4G**). Congruent with increase gastrocnemius mass, MCK-PGC1α had significantly larger mean myofiber CSA than their WT littermates (**Figure 4H**). Muscle power was modestly higher across the 6-min test, however the percent decline in initial power was similar between the genotypes (**Figure 4I**). Quantification of muscular work revealed that MCK-PGC1α mice with HLI performed significantly more work across the 6-min test when compared to WT mice with HLI (**Figure 4J**). Notably, increased muscular work performed by MCK-PGC1α mice occurred despite similar perfusion flux in the 6-min test (**Figure 4K**), reinforcing the hypothesis that improved oxygen utilization likely drives the higher limb function in mice that overexpress PGC1α following HLI.

**Figure 4.**
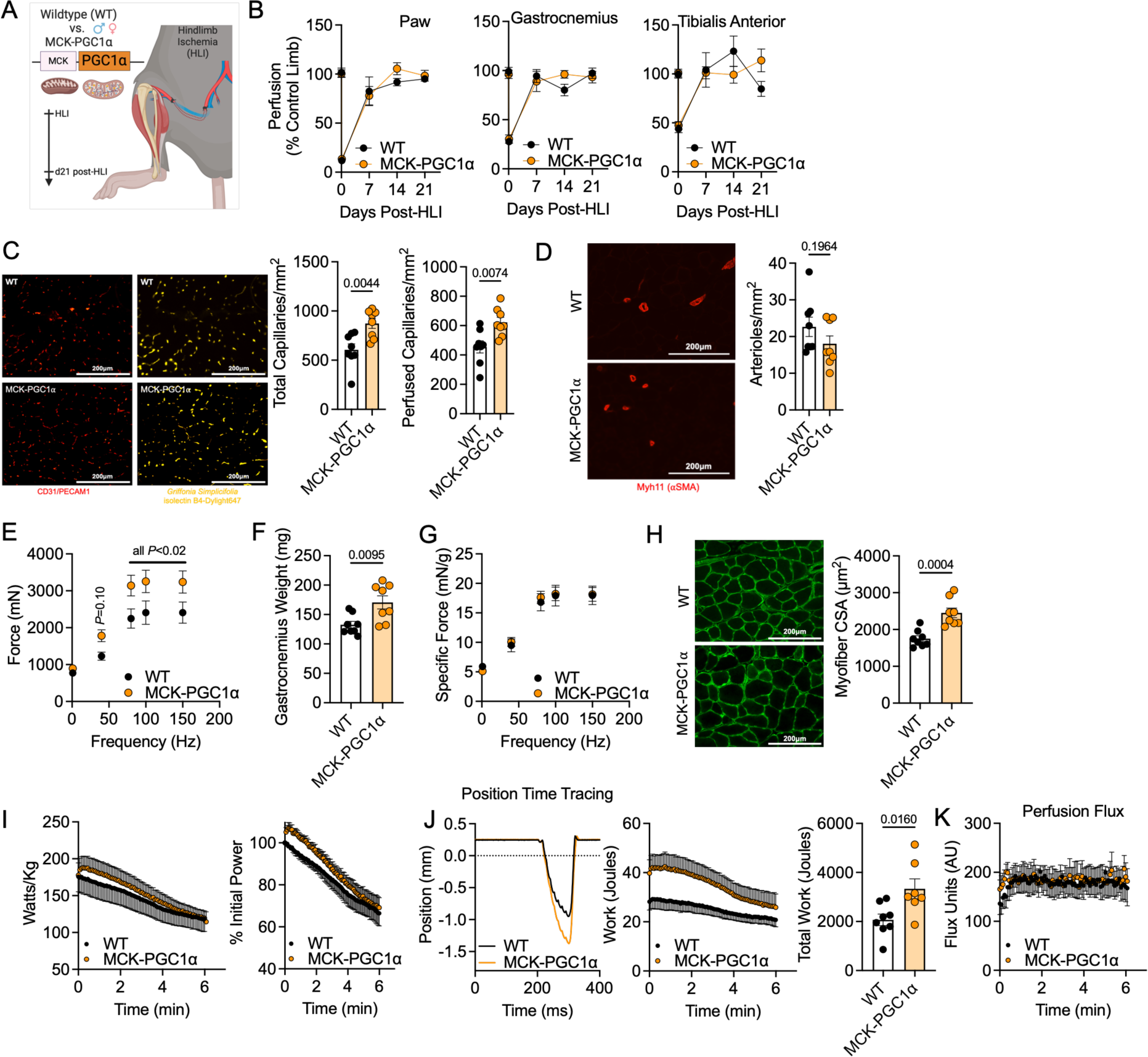
Muscle-specific Overexpression of PGC1α Improves Limb Function Following HLI. (A) Graphic of the experimental design and timeline (generated using BioRender). (B) Laser Doppler flowmetry quantification of perfusion recovery in the paw, gastrocnemius, and tibialis anterior muscles expressed as percentage of control limb (n=10-12/group). Perfusion recovery was analyzed using two-way ANOVA. (C) Representative immunofluorescence images and quantification of total and perfused capillaries. (D) Representative immunofluorescence images and quantification of arterioles. (E) Quantification of absolute forces at different stimulation frequencies. (F) Gastrocnemius muscle mass. (G) Quantification of specific forces at different stimulation frequencies. (H) Representative images and quantification of the myofiber cross-sectional area (CSA). (I) Muscle power across the 6-min test. (J) A representative position-time tracing and work performed across the 6-min test as well as quantification of the total work completed. (K) Perfusion flux during the 6-min test. Panels C-K were analyzed using an unpaired *t*-test (two-tailed). Error bars represent the standard error. All panels contain n=7-8/group.

## DISCUSSION

Preclinical models of hindlimb ischemia (14–21) are the most commonly used approach to study ischemic vasculogenesis, the pathobiology of limb ischemia, and to explore preclinical therapeutics aimed to treat PAD (31, 38–40, 42, 45, 46, 48, 79, 81–96). While there are clear aspects of HLI models that cannot replicate the intricacies of PAD pathogenesis in humans including the chronic nature of atherosclerosis and the diversity of genetics and environmental exposure within the human population, there is a critical need for such models as tools for therapeutic develop which has remained stagnant since the approval of Cilostazol by the Food and Drug Administration (USA) in 1999. In the current work, we sought to develop a functional test that was capable of assessing multiple domains of limb function including muscle blood flow, strength, power, and fatigue. We reasoned that such a test may be better at distinguishing therapeutic benefit than the current ‘gold standard’ of laser Doppler perfusion recovery (22, 97), which is primarily measured in resting conditions where the capacity to increase blood flow to support muscular activity (such as walking) is not assessed. Herein, we present the development and validation of a 6-min limb function test designed for preclinical HLI where the plantarflexor muscle performance and perfusion are assessed using repetitive isotonic (shortening) muscle contractions against a stable load. Using several experimental conditions, we demonstrate the 6-min test can effectively distinguish limb function in the following conditions: 1) inbred mouse strains with divergent responses to HLI; 2) mice with HLI that were randomized to control or exercise therapy groups; and 3) mice subjected to HLI with and without transgenic overexpression of PGC1α, a master regulator of mitochondrial biogenesis and angiogenesis in skeletal muscle.

It has been well characterized that inbred strains of mice and their underlying genetics drive strain-dependent variation in response to HLI (42, 45, 47–50). C57BL/6J mice have more collateral vessel formation (49–51) and prompt a superior recovery in limb perfusion whereas BALB/cJ mice have poor perfusion recovery and are highly susceptible to tissue necrosis (42, 44–49). In the current study, laser Doppler perfusion recovery of both the paw and gastrocnemius muscle were significantly better in C57BL6/J mice compared to BALBc/J mice with HLI (Figure 2A). The corresponding effect sizes (17^2^) were 0.152 and 0.099 for the paw and gastrocnemius, both of which are generally characterized as a large effect. BALBc/J mice also had significantly lower isometric muscle strength (Figure 2F) compared to C57BL6/J mice, with an effect size (17^2^) of 0.67. Using the 6-min limb function test to quantify total muscular work (Figure 2I), the effect size of the strain (17^2^ = 0.74) was substantially larger than that of the laser Doppler perfusion recovery measured in the resting condition. Moreover, laser Doppler analysis of the hyperemic response in C57BL6/J mice compared to BALBc/J during the 6-min limb function (Figure 2J) also had a superior effect size (η^2^ = 0.73) than the gastrocnemius perfusion at rest (17^2^ = 0.099), despite both being statistically significant effects with a *P*<0.05. In totality these results demonstrate the superior ability of this 6-min limb function test to detect strain-dependent differences in limb function following HLI.

Exercise training interventions, including home-based exercise (26, 55), has emerged as one of the most effective treatments to improve or prevent a decline in walking performance in patients with PAD (1, 13, 23, 52–54). Consistent with this body of clinical literature, various forms of exercise training interventions have shown to enhance recovery from HLI in animals (98–100). In the current study, C57BL6/J mice with HLI that were randomized to voluntary wheel running exercise had no improvements in laser Doppler perfusion recovery under resting conditions (Figure 3C) when compared to mice housed without running wheels. However, the exercise trained mice had a greater exercise-induced hyperemia response (Figure 3M) compared to the control (no exercise) group demonstrating that contraction-induced increases in oxygen demand can unmask differences in hemodynamics that were not present under resting conditions. This observation is consistent to results reported by Brevetti *et al*. (101) in which muscle contraction was able to uncover differences in hindlimb blood flow in rats with HLI using microspheres. Exercise trained mice with HLI experienced smaller declines in muscle power output (Figure 3K) and performed ∼45% more total work (Figure 3L) than the control mice with HLI which carried a large effect size (17^2^ = 0.16). These findings are consistent with the multitude of positive effectors of exercise training in the skeletal muscle and vasculature (93, 98, 102–108). It should be noted that exercise training and the control group were all from the C57BL6/J genetic strain of mice and thus have a rapid recovery from HLI. Importantly, the 6-min limb function test was successful at detecting differences in muscular performance and the hyperemic capability of exercise trained mice, whereas resting limb perfusion and muscle strength were not able to distinguish the groups. Similar observations have been reported in patients with PAD. For example, basal/resting microvascular blood volume and flow, measured in the gastrocnemius muscle using contrast-enhance ultrasound, were not different between control subjects without PAD and patients with mild-to-moderate PAD, however differences between these groups were revealed by treadmill or calf muscle exercise (109–111). Similar observations have been reported using magnetic resonance imaging to assess calf muscle perfusion at rest and in response to exercise (112, 113).

Compared to isometric muscle contractions which are most frequently used in rodent studies, isotonic/shortening contractions require more energy (ATP) (114–116) and thus invoke a great demand for oxygen delivery. This feature provides a unique opportunity to evaluate two important features of PAD pathobiology that are believed to contribute significantly to walking performance: 1) impaired blood flow/oxygen delivery and 2) impaired oxygen utilization due to compromised mitochondrial function. To determine the effectiveness of enhancing mitochondrial function in mice with HLI, we employed a transgenic mouse line that overexpresses PGC1α, a well-known controller of mitochondrial biogenesis, which has been previously shown to improve recovery from HLI (79, 80). In the current study, overexpression of PGC1α did not impact laser Doppler perfusion recovery under basal/resting conditions but increased total and perfused capillary densities which is consistent with previous reports (79, 80). Unexpectedly, perfusion flux during the 6-min limb function test was also similar between mice with and without overexpression of PGC1α, despite having clear increases in total and perfused capillary densities. The reason for this discrepancy is unknown at this time but could be related to the superficial penetration depth of the laser Doppler probe whereas the perfused capillaries were quantified across the entire gastrocnemius muscle. Notably, muscle-specific expression of PGC1α improved muscle mass, myofiber area, and absolute force, but specific force levels (an indicator of muscle quality) were identical to WT littermates with HLI (Figure 4). These findings in HLI are consistent with the protective effects of ectopic expression of PGC1α in neuromuscular disease (77, 117), mitochondrial disease (118), and denervation-induced muscle atrophy (78).

This study is not without limitations. First, the acute nature of HLI does not model all aspects of the human PAD condition which is a result of progressive atherosclerotic cardiovascular disease. However, the consistent and easily replicated induction of limb ischemia in the preclinical setting facilitates early testing for novel therapeutics. Second, while there are several advantages of the 6-min limb function test, the fact that it must be performed in anesthetized animals abolishes any impact of pain or neuropathy (i.e., claudication symptoms) on the test performance. This certainly distinguishes the current test from the 6-min walk test performed in most studies involving patients with PAD. The optimal comparison of fatigue properties measured by the decline in peak power would be under contractions with matched workload. While theoretically possible to implement in our conceptual design, it would require different equipment and pose a challenge that hinder implementation of the test. However, the most important aspect for clinical and translational relevance is the work performed during the 6-minute test, which distinguished the effects of disease and therapeutic interventions on muscular performance. In the current study, laser Doppler flowmetry was employed to measure muscle perfusion during the 6-min test. Several other technologies are available to measure muscle blood flow including microspheres (119), microdialysis (120), and indicator dilution methods (121). Other approaches including intravital microscopy and phosphorescence quenching have been used to measure changes in red blood cell flux and microvascular pressure of oxygen (PO_2_) during contractions (122, 123), whereas near infrared spectroscopy could be used to monitor tissue oxygen saturation during this test (124, 125). Leveraging these additional technologies could provide a more comprehensive analysis of convective oxygen delivery, transport, and utilization and the impact on muscle performance in mice with HLI.

In this study, we present the development of a 6-min limb function test that incorporates repeated isotonic contractions and laser Doppler flowmetry to assess hindlimb muscular performance and hyperemia in mice with experimental PAD. This test was successful at detecting differences in ischemic limb function between mouse strains with different severities of HLI pathology, mice with and without exercise therapy following HLI, and those with and without muscle-specific overexpression of PGC1α. In some of these experimental conditions, commonly used outcome measures such as resting/basal perfusion recovery, capillary density, and/or muscle strength were not able to distinguish the groups. Based on this, we propose that the 6-min limb function test may serve as a more robust tool for evaluating potential therapeutics for the treatment of PAD in the preclinical setting.

## Acknowledgements

This study was supported by National Institutes of Health (NIH) grants R01-HL149704 and R01-HL171050 (T.E.R.). L.F.F. was supported by NIH grants R01-HL130318 and R21-AG073239. V.R.P. was supported by a predoctoral fellowship from the American Heart Association (24PRE1193999). A.L.T. was supported by a postdoctoral fellowship from the American Heart Association (24POST1197078). The content is solely the responsibility of the authors and does not necessarily represent the official views of the National Institutes of Health or the American Heart Association.

## Author Contributions

LFF and TER conceived the study and designed the experiments. VRP, JT, ALT, and TER performed experiments and collected data. VRP, JT, ALT, LFF, and TER analyzed and/or interpreted the data. VRP and TER drafted the manuscript. All authors edited, revised, and approved the final version of the manuscript.

